# Dynamic effects of extrinsic noise in a simple oscillatory gene network with delayed negative-feedback regulation: an electronic modeling approach

**DOI:** 10.1101/019638

**Authors:** Moisés Santillán

**Affiliations:** Centro de Investigación y de Estudios Avanzados del IPN, Unidad Monterrey, 66600 Apodaca NL, México

## Abstract

Gene expression is intrinsically stochastic due to the small number of molecules involved in some of the underlying biochemical reactions. The resulting molecule-count random fluctuations are known as biochemical noise. The dynamic effects of intrinsic noise (that originated within the system) have been widely studied. However, the effects of the noise coming from other sources the system is in contact with, or extrinsic noise, is not so well understood. In this work we introduce an electronic model for a simple gene oscillatory network, with delayed negative-feedback regulation. Notably, this model accounts for the intrinsic biochemical noise due to the slow promoter switching between the active and inactive states; but dismisses biochemical noise due to mRNA and protein production and degradation. We characterize the oscillatory behavior of this gene network by varying all the relevant parameter values within biologically meaningful ranges. Finally, we investigate how different sources of extrinsic noise affect the system dynamic behavior. To simulate extrinsic noise we consider stochastic time series coming from another circuit simulating a gene network. Our results indicate that, depending on the parameter affected by extrinsic noise and the power spectra of the stochastic time series, the system quasi-periodic behavior is affected in different ways.

## I. INTRODUCTION

Periodic phenomena are ubiquitous in biology [1]. They can be observed from the single cell to the ecological levels. Genetic oscillators are of particular interest because they are the basis of biological phenomena like the circadian clock [2] and the segmentation clock of somitogenesis [3].

Genetic oscillators have also been a subject of interest in synthetic biology [4, 5]. However, the extremely regular behavior of natural genetic oscillators contrasts with the irregularity of oscillations observed in most synthetic oscillators. From a theoretical point of view, the fact that synthetic genetic oscillators are irregular is not surprising because of the ever present biochemical noise [6, 7].

Given that some of the molecules involved in gene expression and gene regulation occur in low numbers (less than ten in some cases), the stochastic nature of the corresponding biochemical reactions becomes evident. Thus, the plots of molecular counts vs. time present stochastic fluctuations around the curves predicted by the corresponding deterministic descriptions, and these fluctuations are known as biochemical noise [6, 7]. In particular, if a deterministic description of the system predicts that a particular molecule count oscillates periodically in time, biochemical noise will make the oscillation period and amplitude irregular. Typically, these irregularities increase as the noise amplitude increases, to the point that the system periodic behavior can get masked. The question is then, how come natural genetic oscillators can be so regular despite biochemical noise. To the best of our knowledge, this is still an open question.

For a given system, biochemical noise can be either intrinsic or extrinsic. Noise originated by biochemical reactions within the system is termed intrinsic noise. On the contrary, biochemical noise affecting the system dynamics but originated within other systems is known as extrinsic noise. In a recent work [8], the authors proved that, depending on their time scale, extrinsic fluctuations can affect the performance of biochemical networks in many different ways. This suggests that interaction mechanisms between gene regulatory networks may have been fine tunned by evolution in such a way that the associated extrinsic noise improves, or at least does no affect, the gene networks’ performance. Taking this into consideration, it is interesting to study how extrinsic noise with different characteristics affects the performance of genetic oscillators.

Inspired on the previous paragraphs discussion, the present work is advocated to studying the influence of different sources of extrinsic noise on the dynamics of a simple gene regulatory network, known to have a periodic behavior: a single gene subject to delayed negative-feedback regulation. To achieve this objective, we designed and built an electronic circuit that simulates the gene network stochastic behavior, and studied its dynamics. To account for extrinsic noise, we interconnected in different ways two of such circuits, assuming that one of them is the source of extrinsic noise for the other. Given that we make use of both analog and digital electronics to mimic the dynamic behavior of a natural phenomena, the present work is related to the long tradition of analog and hybrid computing [18].

## II. MODEL DEVELOPMENT

### A. Gene Regulatory Network

Consider the hypothetic gene regulatory pathway schematically represented in Fig. 1 and described in detail below. When active, a promoter is transcribed, and so mRNA molecules are synthesized, at a rate *k*_*M*_. The resulting mRNA molecules are randomly degraded, with a degradation rate constant *γ*_*M*_. Before being degraded, mRNA molecules are translated and proteins are synthesized at a rate *k*_*P*_. The translated proteins catalyze the synthesis of metabolites *W*, and they are randomly degraded with a degradation rate constant *γ*_*P*_. Furthermore, metabolites *W* bind activator molecules *A*, to inhibit them. Uninhibited activators can bind an inactive promoter, activating it. When the activator detaches from the promoter, it turns inactive.

**FIG. 1:**
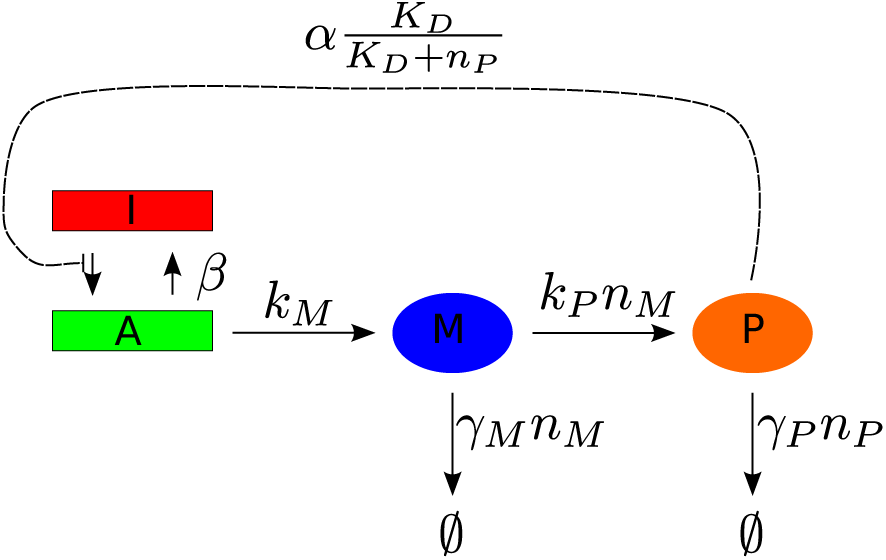
Schematic representation of the gene regulatory network here studied.

Under the assumption that the metabolite production rate is proportional to the protein molecule count, and that metabolite degradation occurs randomly, the differential equation governing metabolite dynamics is

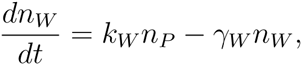

where *n*_*P*_ and *n*_*W*_ respectively denote the protein and metabolite molecule counts, *k*_*W*_ is the metabolite synthesis rate per protein, and *γ*_*W*_ is the metabolite degradation rate constant. If the metabolite synthesis and degradation processes are much faster than the mRNA and protein dynamics, a quasi-steady state approximation can be made for the metabolite dynamics (*dn*_*W*_ */dt* = 0), and so the number of metabolites is given in terms of the protein count by the following expression:

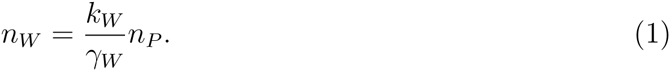

As previously asserted, we assume that activator molecules are inhibited when they are bound by *W* metabolites and that the corresponding chemical reaction is

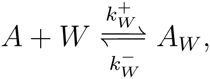

in which *A* and *A*_*W*_ respectively represent uninhibited and inhibited activators, while 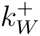 and 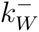 are the forward and backward reaction rate constants. Under the supposition that this reaction rapidly equilibrates with the rest of the system dynamics, and assuming that the total activator count is constant,

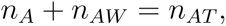

the number of uninhibited activators comes out to be

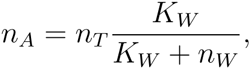

with 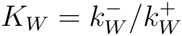. By substituting Eq. (1) into this last expression we obtain

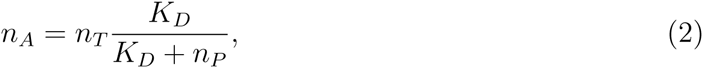

where *K*_*D*_ = *K*_*WW*_=*k*_*W*_.

Let us represent the promoter activation-inactivation process by means of the following reaction

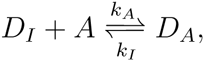

where *D*_*I*_ and *D*_*A*_ respectively represent inactive and active promoters, while *k*_*A*_ and *k*_*I*_ are the activation and deactivation reaction rate constants. Recall that *A* denotes an uninhibited activator. The corresponding forward and backward reaction rates are respectively given by

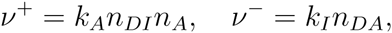

where *n*_*DI*_ and *n*_*DA*_ respectively represent the numbers of inactive and active promoters. If we now substitute Eq. (2), the forward reaction rate transforms into

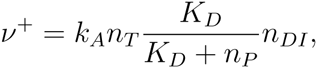

Hence, we can alternatively represent the promoter activation-inactivation process by means of the following reaction

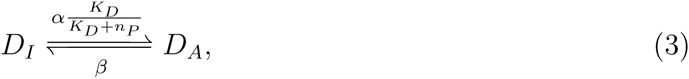

with

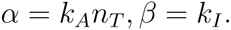

From the above discussion we can see that gene expression can be modeled in general as a set of chemical reactions. On the other hand, chemical reactions are intrinsically stochastic and their stochastic nature becomes apparent in the low molecular-count limit. Since this is the case for many of the reactions involved in gene expression, the concomitant stochastic fluctuations (also known as intrinsic noise) cannot be ignored. Several works have studied the consequences of having stochastic synthesis and degradation of mRNA and protein molecules. Interestingly, only a few of them have dealt with the stochasticity of promoter gating between the active and inactive states [9]. In the present work we are interested in studying the dynamics of a gene regulatory network with negative feedback, in which promoter gating is the only source of intrinsic noise, and in which the time delays due to transcription and translation are accounted for. Therefore, the previously discussed gene regulatory scheme can be modeled as follows.

A telegraph stochastic process can be employed to simulate promoter dynamics, taking into consideration that the transition probabilities per unit time (propensities) between the inactive and active states are respectively given by:

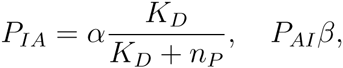

where *P*_*IA*_ and *P*_*AI*_ respectively denote the propensities for the transitions from and inactive to the active, and from the active to the inactive states. Furthermore, since we decided to disregard the intrinsic noise associated to the mRNA and protein dynamics, they can be modeled by means of the following differential equations:

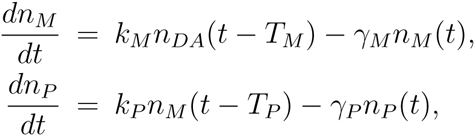

where *T*_*M*_ and *T*_*P*_ are the time delays due to transcription and translation.

As previously discussed, the model just developed corresponds to a single gene network subject to delayed negatve-feedback regulation. This is a very simple system which, due to its simplicity can be thoroughly analyzed. On the other hand, despite its simplicity, the present model is biologically sound because it shares the architecture of the gene network behind the segmentation clock of somitogenesis [3].

### B. Electronic analog for the gene network

As we have seen, the equations governing the mRNA and protein dynamics are of the form

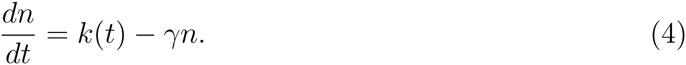

where *n* represents the molecule count, *k*(*t*) is the molecule synthesis rate, and *γ* is the corresponding degradation rate constant. Let us assume that function *k*(*t*) is upper-bounded: *k*(*t*) *≤ k*_*max*_. Then, by defining the dimensionless variable

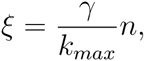

Eq. (4) can be rewritten as follows:

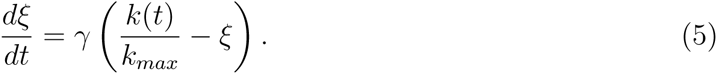

Consider now the RC circuit depicted in Fig. 2. If the ground voltage is set equal to 0 V, one can easily prove—by equating the currents through the resistance and the capacitor — that voltage *V*_*C*_ satisfies the following differential equation,

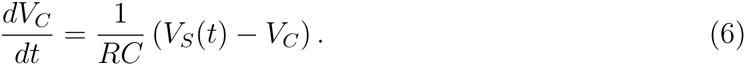

**FIG. 2:**
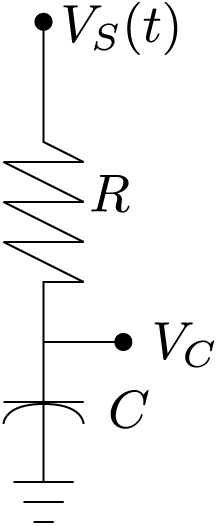
Schematic representation of a typical RC circuit

Assume that *V*_*S*_(*t*) is upper bounded: *V*_*S*_(*t*) *≤ V*_*max*_ and define *ζ* = *V*_*C*_*/V*_*max*_. Then, from Eq. (6), variable *ζ* satisfies the following differential equation:

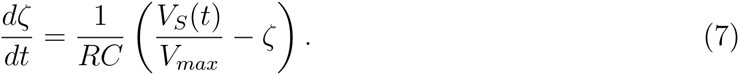

Note that Eqs. (5) and (7) are identical, allowing us to conclude that an RC circuit can be used to model the dynamics of molecule synthesis and degradation, whenever the intrinsic noise inherent to molecule synthesis and degradation can be disregarded. Observe that the time constant *RC* is analog to the inverse of the degradation rate constant *γ*, while the normalized voltage *V*_*S*_(*t*)*/V*_*max*_ is analog to the normalized synthesis rate *k/k*_*max*_.

From the discussion in the former paragraphs, the gene regulatory network schematically represented in Fig. 1 can be modeled by the electronic circuit represented in Fig. 3.

**FIG. 3:**
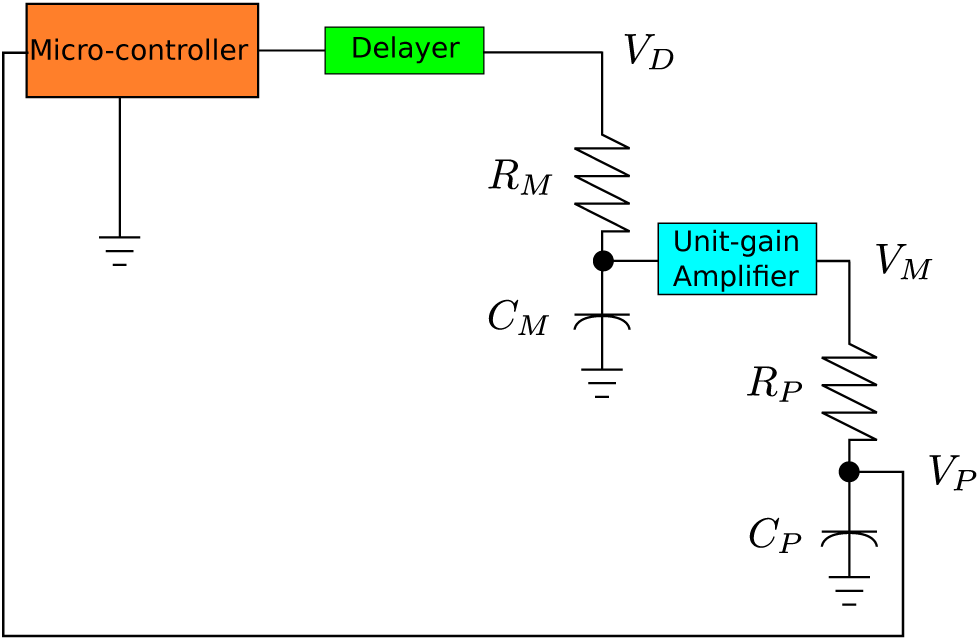
Schematic representation of an electronic circuit that models the gene regulatory network in Fig. 1. See the main text for details.

The different circuit elements depicted in Fig. 3 play the following roles:

- Resistance *R*_*M*_ and capacitor *C*_*M*_ account for the dynamics of mRNA molecules. The mRNA degradation rate is given by 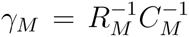, while the voltage *V*_*D*_(*t*) is proportional to the mRNA transcription initiation rate. In accordance to the promoter dynamics, *V*_*D*_(*t*) randomly shifts between zero (inactive Promoter) and the maximum (active promoter) voltage value. The corresponding propensities (transition probabilities per unit time) are those of promoter activation and deactivation.
- Resistance *R*_*P*_ and capacitor *C*_*P*_ account for the dynamics of proteins. The protein degradation rate is given by 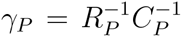, while the voltage *V*_*M*_ (*t*) is proportional to both the mRNA molecule count and the translation initiation rate. On the other hand, voltage *V*_*P*_ (*t*) is proportional to the protein count.
- The unit-gain amplifier separating the mRNA and protein RC circuits ensures that the protein RC circuit does not affect the dynamics of the mRNA RC circuit.
- The micro-controller plays the role of the promoter. It employs one pin to measure voltage *V*_*P*_ (*t*) and uses this information to randomly compute the voltage (either zero or high) at the output pin. Below we describe in detail the algorithm the micro-controller uses to perform this task.
- In order to account for the delays due to transcription, translation, post-translational protein modifications, etc., we include a shift-register integrated circuit (IC) between the micro-controller and the mRNA RC circuit. This IC delays the digital signal coming out from the micro-controller a given amount of time (the corresponding mechanisms will be explained below). As previously asserted, time delays originate at various stages in a real gene network. However, we are interested in the effects these delays have on the negative feedback regulatory loops. It can be demonstrated that, concerning the fixed point stability, since the various delays occur in sequential processes, they can be lumped together in a single delay taking place anywhere in the loop [10]. Concomitantly, we decided to include a single lumped delay at the level of transcription.

Before analyzing the micro-controller algorithm, consider that this device performs iteratively, repeating its program in short duration loops. In our case, we decided to employ the popular Arduino Uno [19]. According to our measurements, this micro-controller loops last 0.28 ms in average. Taking all this into account, the micro-controller algorithm is as follows:

1. Voltage *V*_*P*_ is measured at the input pin.
2. The probabilities of flipping from the active to the inactive (*P*_*AI*_), and from the inactive to the active (*P*_*IA*_) states in a micro-controller cycle are respectively computed as *P*_*AI*_ = *β* and *P*_*IA*_ = *α * K*_*D*_/(*K*_*D*_ + *V*_*P*_), where *β*, *α*, and *K*_*D*_ are parameters whose values are estimated below.
3. A random number *r* is generated from a uniform random distribution to determine whether the promoter changes its state, given the previously computed transition probabilities.
4. If the promoter is active and *r ≤ P*_*AI*_ the promoter state is changed to inactive and the voltage at the output pin is set to 0 V. Otherwise, the promoter remains active and the output voltage is set to its maximum value (5 V in our case).
5. Conversely, if the promoter is inactive and *r ≤ P*_*IA*_, the promoter is set to active and the output voltage is set to its maximum value. Otherwise, the promoter remains inactive and the output voltage is set to 0 V.
6. The voltage at a given digital output pin, that sends a clock pulse to the shift register (the delayer), is set to its maximum value and immediately it is set back to 0 V.

The shift register (SR) consists of a series of memory devices that can be set to either 0 V or the maximum voltage (which in this case is 5 V). Each time the SR gets a clock pulse, every one of its memory devices assumes the state of the previous device, and the first memory device assumes the value of the input voltage. In our case, this input voltage corresponds to state of the promoter. Given that a clock pulse arrives each time the micro-controller completes a cycle, the voltage corresponding to the current promoter state reaches the last SR memory device after *n* micro-controller cycles, were *n* is the number of SR memory devices. Thus, by modifying the value of *n* one can control the time delay between the change of state at the micro-controller output pin and the moment this change is felt by the RC circuit corresponding to the mRNA dynamics.

### C. Parameter values

As previously mentioned, the average cycle duration of the micro-controller is

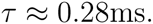

This is important because, from the way the electronic circuit is built, this parameter plays the role of time unit for the whole system.

The maximum probability rates from the inactive to the active, and from the active to the inactive states are respectively given by:

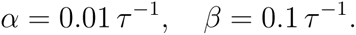

This values imply that, in the long term, the promoter spends only about 10% of the time in the active state, and that, in the absence of regulation, the average promoter residence times in the active the inactive states are 10 and 100 microcontroler cycles, respectively.

The resistance and the capacitor corresponding to the mRNA dynamics have the following values:

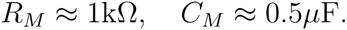

This corresponds to a degradation rate constant of

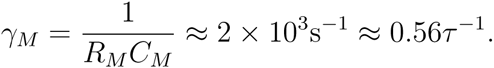

The resistance and capacitor values corresponding to the protein dynamics are

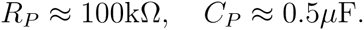

This implies a degradation rate constant of

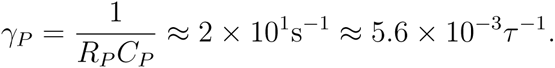

After implementing the electronic circuit with the above parameter values, with no time delay and with no feedback (*K*_*D*_ *→ ∞*), we measured the voltage time series corresponding to protein dynamics and computed an average value of 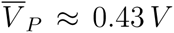. We are interested in sustained oscillations generated by a delayed negative feedback, and it is known that a strong feedback loop (small *K*_*D*_ value) is necessary for that purpose [11–13]. Therefore, we chose the following values for parameter *K*_*D*_:

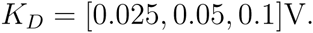

Finally, large delay values are necessary for an oscillatory behavior [11–13]. Thus, taking into consideration that the protein half life is about 125*τ*, we assumed the following time delay values

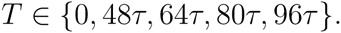

These parameter values guaranty the following behavior.

- The promoter becomes active sporadically and it remains active for a short time (a few times *τ*), in agreement with the experimentally observed transcriptional bursting [9, 14].
- mRNA molecules have a half-life (*t*_1/2_ = ln 2*/γ*_*M*_ *≈* 1.23*τ*) comparable to the system time unit. It has been observed in vivo that mRNA molecules have short half lives in general, but in particular their half life is of the order of a minute for some prokaryotic genes [15].
- The protein degradation rate is about 100 times smaller than the mRNA degradation rate, in agreement with the fact that, in general, proteins have much longer half-lives than mRNA molecules [15].
- The time delay and the value of parameter *K*_*D*_ were chosen so that the system shows an oscillatory behavior. It has been reported in deterministic systems that *K*_*D*_ needs to be small, as compared with the maximum possible molecule count, and that the time delays has to be long, in terms of the mRNA half life, in order to observe sustained oscillations [11–13]. Hence, taking this into consideration, assumed that *T* values are of the order of the protein half life, and took small *K*_*D*_ values as compared with the average protein voltage in the open loop configuration (see the Results for further details).

## III. RESULTS

### A. Characterization of system dynamic behavior

Data, in the form of voltage time series measured at point *V*_*P*_ of the circuit (see Fig. were acquired by means of a 

~~~
BitScope
~~~

 BS10 (manufactured by BitScope[20]), controlled from 

~~~
Python 2.7
~~~

 via the public library 

~~~
BitScope
~~~

 Library 2.0. Recall that, according to our model, voltage *V*_*P*_ reports the amount of protein, so the recorded time series allow us to see how the protein level evolves in time. Unless stated otherwise, we employed a sampling frequency of 500 samples per second which, as we shall see, is more than enough for the purposes of the present work. In Fig. 4 we plot two examples of such time series, which correspond to a network with no feedback (or open loop, *K*_*D*_ *→ ∞*), and to a network with delayed negative feedback in which the relevant parameter values are *K*_*D*_ = 0.05 V and *T* = 80*τ.*

**FIG. 4:**
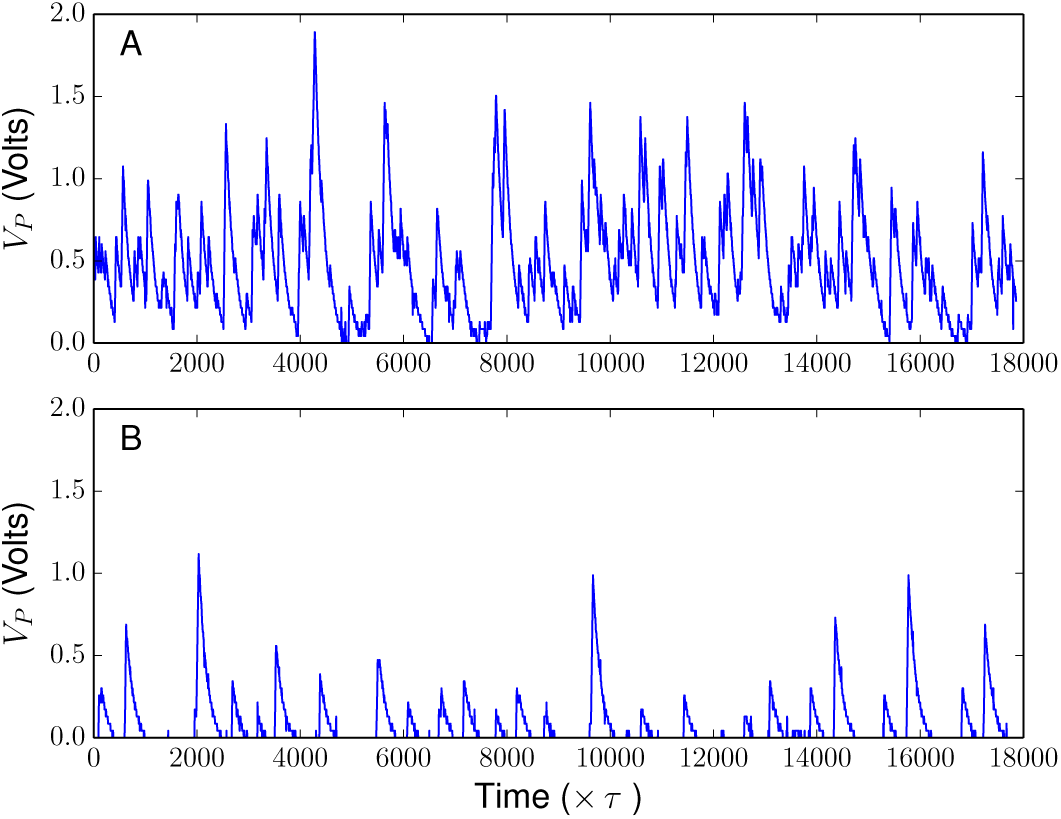
Recordings of the voltage reporting protein levels in the circuit designed to simulate a simple gene network with delayed negative feedback. A) Gene network without feedback (or open loop, *K*_*D*_ *→ ∞*). B) Gene network with *K*_*D*_ = 0.05 V and *T* = 80*τ*.

As previously discussed, we expect the negative feedback and the time delay to induce some sort of oscillatory behavior. Indeed, both time series in Fig. 4 are notoriously different, and some regularity can be observed in the delayed-feedback case. However, at first sight it is impossible to judge whether the system behaves periodically or not. To elucidate this point and to further understand the influence of parameters *K*_*D*_ and *T*, we carried out a Fourier analysis as follows. For a given set of *K*_*D*_ and *T* parameter values, we recorded 1,000 independent time series with a rate of 500 samples per second, each time series being 10,000 data points long. Then, we employed the algorithm 

~~~
numpy.rfft
~~~

 of 

~~~
python
~~~

 to compute the fast Fourier transform of each time series, and used the result to calculate the corresponding power spectrum. Finally, we obtained the average power spectrum (averaged over the 1,000 measured time series) and plotted it. The whole procedure was repeated for different combinations of parameters *K*_*D*_ and *T*, and the results are shown in Fig. 5.

**FIG. 5:**
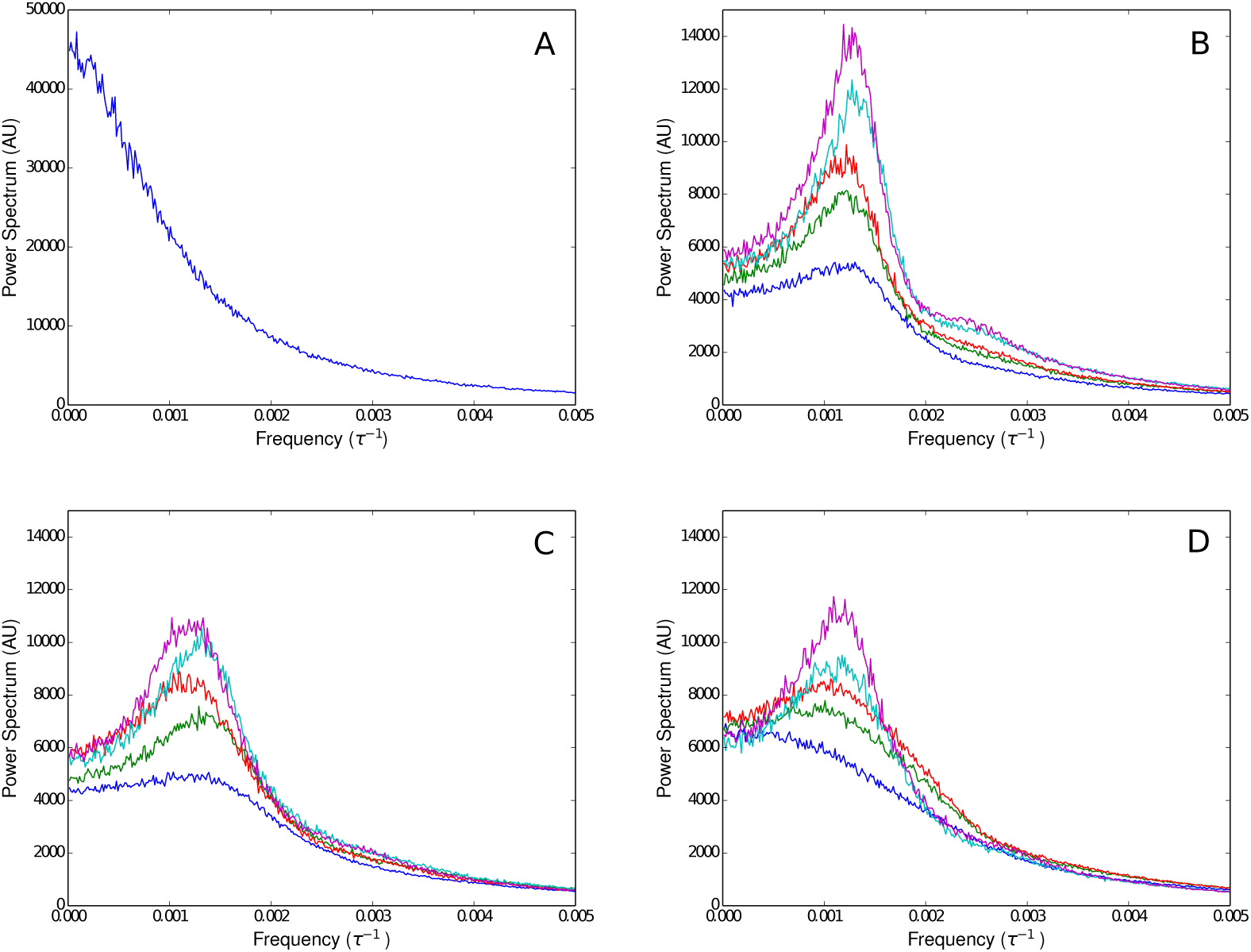
Power spectra computed from the recorded *V*_*P*_ time series, with different combinations of parameters *K*_*D*_ and *T*. A) Open loop configuration, *K*_*D*_ *→ ∞*, B) *K*_*D*_ = 0.025 V, C) *K*_*D*_ = 0.05 V, and D) *K*_*D*_ = 0.1 V. The color code for the plots in graphs B-D is a follows: blue, *T* = 0; green, *T* = 48*τ*; red, *T* = 64*τ*; cyan, *T* = 80*τ*; magenta, *T* = 96*τ*.

Observe that, in the open-loop configuration, the power spectrum is monotonically decaying and thus corresponds to colored noise. This further implies that, in this case, the electronic circuit has no oscillatory behavior. Regarding the behavior or the gene circuit with delayed negative-feedback regulation, we see that the power spectrum is in most cases a concave function with a single maximum. This type of power spectrum is indicative of a quasi-periodic behavior. Furthermore, quasi-periodicity is not apparent when the time delay is very small and parameter *K*_*D*_ attains large values (*K*_*D*_ *≈* 0.1 V). But for instance, when *K*_*D*_ = 0.025 V a quasi-oscillatory behavior is observed even for *T* = 0. Concerning the influence of parameters *K*_*D*_ and *T* on the system dynamics, we can see in Fig. 5 that decreasing the value of *K*_*D*_ and/or increasing that of *T* increases the height of the power-spectrum maximum point. This is in agreement with the notion that increasing the time delay and/or the strength of the feedback increases the amplitude of periodic oscillations. Interestingly, the position of the spectrum maximum point remains unchanged despite variations in *K*_*D*_ and *T*. This finding contradicts what is normally observed in deterministic systems in which the frequency of oscillations usually decreases as the time delay increases [16].

To finish the system characterization, we repeated the previously described experiments with different values of mRNA and protein degradations rates. In particular, we halved and doubled both degradation rates by accordingly modifying resistances *R*_*M*_ and *R*_*P*_. The resulting power spectra are shown in Fig. 6. Observe that changing mRNA degradation rate has no notorious effect upon the system oscillatory behavior. However, modifying protein degradation has important consequences. Increasing the protein degradation rate (*γ*_*P*_) makes the spectra tails heavier in both the open loop and the delayed negative feedback configurations. This means that higher frequency components contribute more as *γ*_*P*_ increases. In particular, in the delayed negative feedback configuration, the maximum point of the power spectrum achieves larger values and moves to higher frequencies as the protein degradation rate increases.

**FIG. 6:**
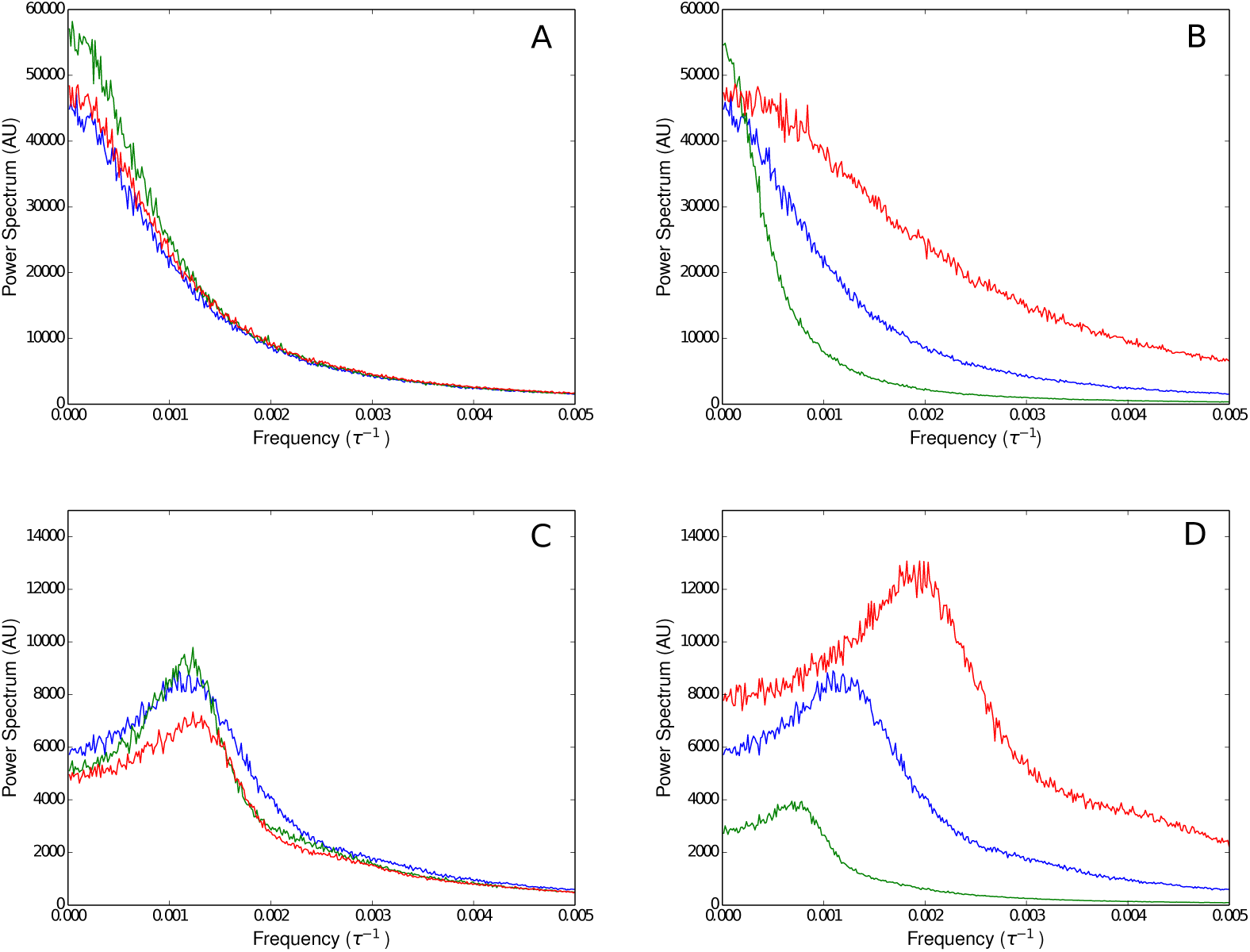
Power spectra computed from the recorded *V*_*P*_ time series, with different values of the mRNA and protein degradation rates. Panels A and B correspond to an open-loop gene circuit, while panels C and D correspond to a gene circuit with negative feedback (*K*_*D*_ = 0.05 V) and time delay (*T* = 64*τ*). In panels A and C we can see the effect of changing the mRNA degradation rate; line color code is as follows: blue corresponds to the nominal parameter values, green corresponds to a halved mRNA degradation rate, and red corresponds to a doubled mRNA degradation rate. Finally, in panels B and D we can see the effect of changing the protein degradation rate; line color code is as follows: blue corresponds to the nominal parameter values, green corresponds to a halved protein degradation rate, and red corresponds to a doubled protein degradation rate

### B. Dynamic effects of extrinsic noise

So far, we have investigated the dynamic behavior of a gene network subject to delayed negative feedback regulation, in which the only source of biochemical noise is the promoter random gating between the active and inactive states. Since this kind of noise originates within the system, it is known as intrinsic noise [6, 7, 17]. However, the system can also be affected by extrinsic noise. That is, biochemical noise originated within other systems that interact with the current system [8]. For instance, in our case, we have seen that parameter *K*_*D*_ is proportional to the number of activator molecules. However, these activators are produced via the expression of another gene and so, rather than being constant, *K*_*D*_ can be understood as the realization of a stochastic process.

According to the previous subsection results, the system quasi-periodic oscillatory behavior is quite sensitive to changes of parameters *K*_*D*_ and *γ*. Thus, it is interesting to investigate the influence that random extrinsic-noise variations of these parameters have on the system qausi-periodic behavior. To do this, we built two electronic circuits (termed A and B) like that schematically represented in Fig. 3, and assume that they interact according to one of the following situations:

#### Situation I

Circuit A simulates a gene network that produces the transcription factors (activators) that regulate the expression of the gene network simulated by circuit B.

#### Situation II

Circuit A simulates a gene network that produces the proteases that degrade the proteins produced by the gene network simulated by circuit B.

**FIG. 7:**
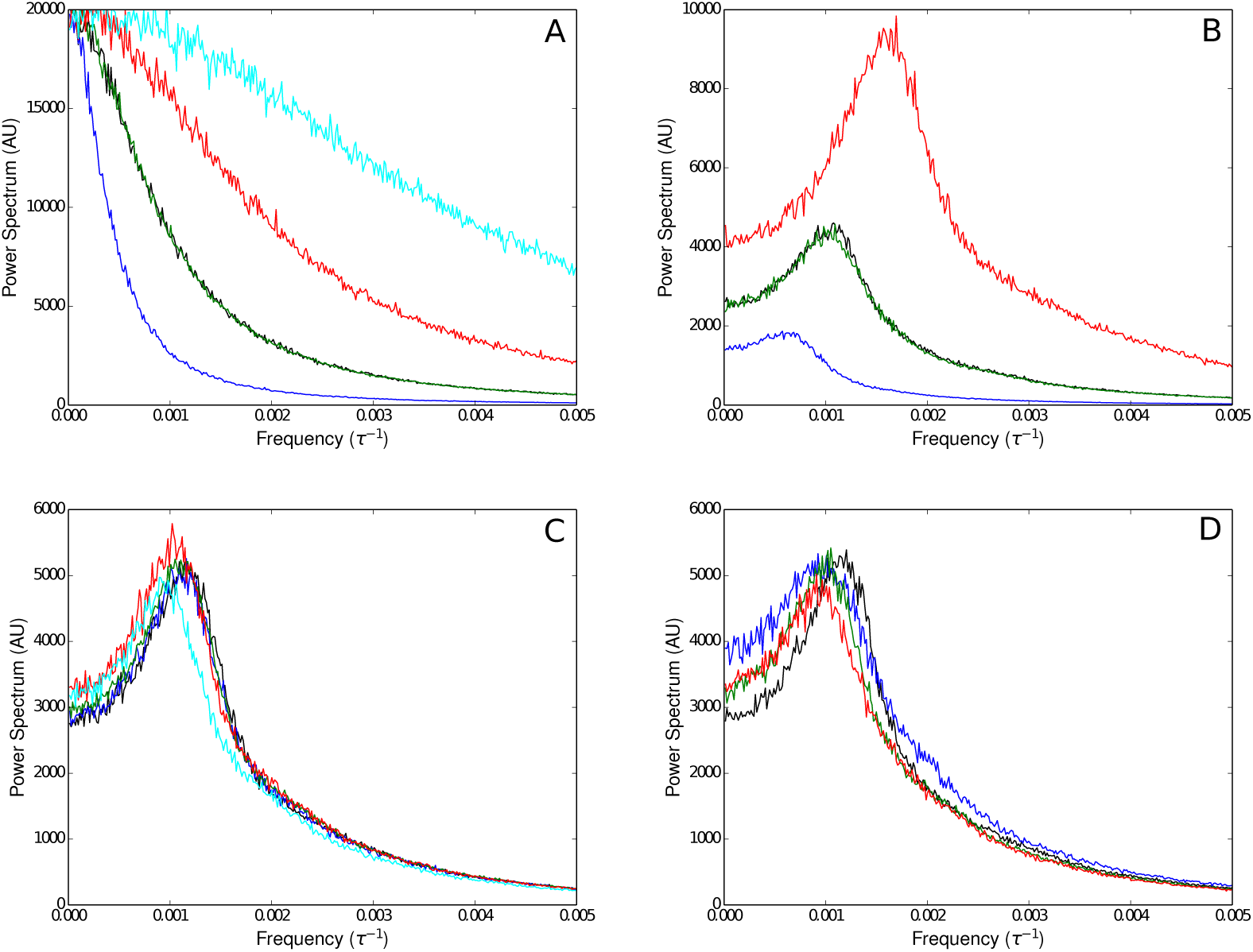
Power spectra computed from the recorded *V*_*P*_ time series of two coupled gene networks: one producing the transcription factors (panels A and B) that regulate the expression of the other gene network (C and D). We considered two different scenarios for the transcription-factor producing gene network: it expresses constitutively (panels A and C), and it is subject to delayed negative-feedback self-regulation with *K*_*D*_ = 0.05 V and *T* = 64*τ* (panels B and D). In each case we considered different transcription-factor (TF) degradation rates, indicated by means of a color code as follows: black corresponds to TF degradation rate equal to that of the regulated-network protein, but with the two networks uncoupled; all other line colors correspond to coupled gene networks; blue corresponds to a TF degradation rate half as large as that of the regulated protein; green corresponds to a TF degradation rate equal to that of the regulated protein; red corresponds to a TF degradation rate twice as large as that of the regulated protein; and magenta corresponds to a TF degradation rate five times as large as that of the regulated protein.

Taking into account that *K*_*D*_ is proportional to the number of activators, we electronically simulated Situation I as follows:

- All parameters of circuit B, with the exception of *K*_*D*_ and *T*, were set to their nominal values. Parameter *T* was set to 64*τ*, and the value of *K*_*D*_ was chosen as explained below.
- One of the analog input pins of the micro-controller of circuit B is employed to measure voltage *V*_*P*_ of circuit A (recall that *V*_*P*_ is proportional to the protein count). Then, circuit B’s *K*_*D*_ value is computed as *K*_*D*_ = *εV*_*P*_, with *ε* chosen so the average value of *K*_*D*_ equals 0.05 V
- Most of the parameters of circuit A (with the exception of *K*_*D*_ and *γ*) are set to their nominal vales. In particular, we set *T* = 64*τ*. Regarding *K*_*D*_ we considered two possible values for it: *K*_*D*_ *→ ∞*, which corresponds to a constitutive-expressing gene, and *K*_*D*_ = 0.05 V, that corresponds to a gene network with delayed negative feedback regulation. Finally, we halved, doubled, and increased to five times the nominal value of parameter *γ*_*P*_, aimed at changing the power spectrum characteristics of circuit A’s *V*_*P*_ time series.

After assembling the two interacting circuits and connecting them, we recorded the *V*_*P*_ voltages of each one of them by means of a bitscope. Each recording was 5,000 data points long, and was performed with a frequency of 250 samples per second. Then, we computed the power spectra of the time series recorded from both circuits. The whole procedure was repeated 1,000 times, and the average power spectra (averaged over the 1,000 experiment repetitions) were computed at the end. The results are shown in Fig. 8.

**FIG. 8:**
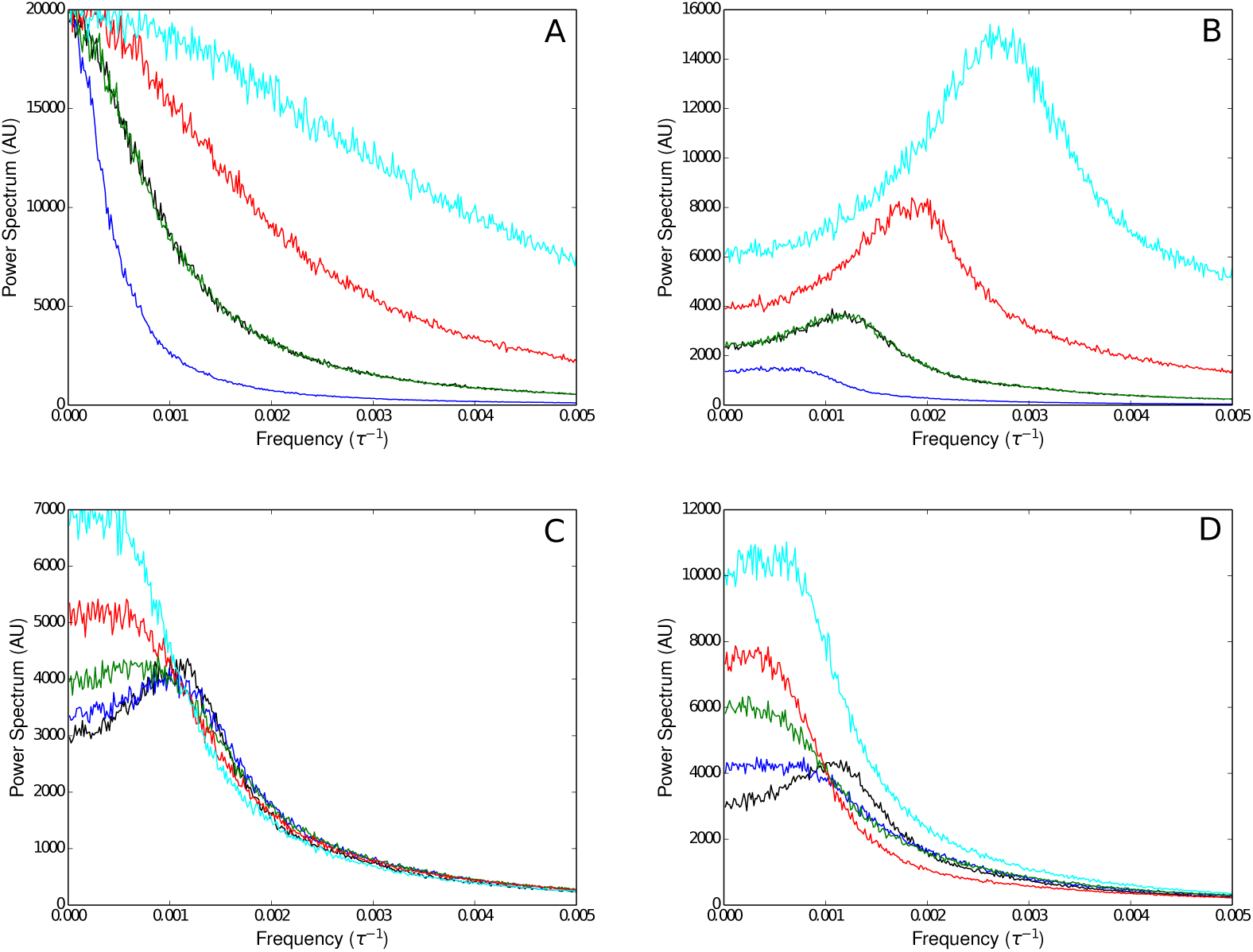
Power spectra computed from the recorded *V*_*P*_ time series of two coupled gene networks: one producing the proteases (panels A and B) that degrade the proteins resulting from the expression of the other gene network (C and D). We considered two different scenarios for the protease producing gene network: it expresses constitutively (panels A and C), and it is subject to delayed negative-feedback self-regulation with *K*_*D*_ = 0.05 V and *T* = 64*τ* (panels B and D). In each case we considered different protease degradation rates, indicated by means of a color code as follows: black corresponds to protease degradation rate equal to that of the target protein, but with the two networks uncoupled; all other line colors correspond to coupled gene networks; blue corresponds to a protease degradation rate half as large as that of the degraded protein; green corresponds to a protease degradation rate equal to that of the degraded protein; red corresponds to a protease degradation rate twice as large as that of the degraded protein; and magenta corresponds to a protease degradation rate five times as large as that of the degraded protein.

From Figs. 8A and 8C we can conclude that colored-noise random variations in the transcription factor count have little effect upon the quasi-periodic oscillatory dynamics of the regulated gene network. The most noticeable difference, as compared with the case with no extrinsic noise, is observed when the transcription factor degradation rate if five times larger that that of circuit B’s proteins. Since in this last case the transcription-factor power spectrum has larger high-frequency contributions, our results may indicate that highfrequency random fluctuations in the transcription factor count have more important effects upon the quasi-periodic oscillatory behavior of circuit B.

Figs. 8B and 8D also reveal that the quasi-periodic oscillatory behavior of circuit B is extremely robust to quasi-periodic random fluctuations of the transcription factor count. Given that the transcription-factor power spectra are very different, it is not possible to analyze the results in terms effects due to low-frequency and high-frequency extrinsic-noise contributions.

To simulate Situation II, consider that in our circuits, protein degradation rate is inversely proportional to resistance *R*_*P*_. On the other hand, if circuit A produces the proteases that degrade the proteins of circuit B, one could expect that circuit’s B protein degradation rate is proportional to voltage *V*_*P*_ of circuit A. Thus, Situation II can be implemented by controlling circuit’s B *R*_*P*_ resistance with circuit’s A *V*_*P*_ voltage. We achieved this by using digital potentiometer DS1803 (1,000 kΩ maximum resistance) controlled by an arduino; and by programming the arduino such that it reads voltage *V*_*P*_ in circuit A and sets resistance *R*_*P*_ of circuit B to *ξ/V*_*P*_. Parameter *ξ* was chosen so that 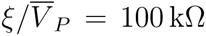, with 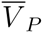 the average value of circuit A *V*_*P*_ voltage.

After implementing the electronic circuit that simulates Situation II, we repeated the experiments of Situation I and report the results in Fig. 8. Observe that no matter whether the protease gene expresses constitutively or its expression level shows random quasi-periodic oscillations, this kind of extrinsic noise always has a deleterious effect upon circuit B quasi-periodic behavior. Moreover, the effects of this extrinsic noise augment as the protease degradation rate increases and the hight frequency contributions to the corresponding power spectra become greater.

## IV. CONCLUDING REMARKS

An electronic circuit that simulates the dynamic behavior of a gene network subject to delayed negative-feedback regulation was designed and built. Notably, this circuit assumes that the only source of intrinsic noise in the regulatory pathway is the promoter stochastic gating between the active and inactive states. Regarding the model parameter values, they were chosen so they fall within biologically plausible ranges, as well as to ensure that the circuit shows a quasi-periodic dynamic behavior.

Before investigating the effects of extrinsic noise on the system dynamics, we investigated the robustness of its dynamic behavior to variations of the model parameters. To that end, we centered our attention on the power spectrum of the recorded *V*_*P*_ time series, where *V*_*P*_ is the circuit voltage that is analog to the concentration of proteins in the gene regulatory network.

First of all we proved that, as expected, the built circuit displays a quasi-periodic behavior when it is subject to delayed negative-feedback regulation, and that this behavior dissappears in the absence of a feedback regulatory loop. Furthermore, we were able to show that either increasing the feedback-loop time delay or strengthening the feedback-loop enhances the system quasi-periodic behavior, in agreement to what happens in the correspondig deterministic systems. Finally, we studied how changes of the mRNA and protein degradation rate constants affect the observed quasi-periodic oscillations. We found that the system quasi-periodic behavior is quite robust to mRNA degradatio rate changes, but sensitive to modifications of the protein degradation rate constant, *γ*_*P*_. Larger *γ*_*P*_ values make the amplitude and frequency of oscillations increase.

To simulate extrinsic noise we took stochastic time series coming from another circuit simulating a gene network. Let us term circuit A the circuit where extrinsic noise is originated, and circuit B the one influenced by extrinsic noise. We considered two different scenarios

- Circuit A produces the transcriotion factors (which in this case are activators) that regulate the expression of circuit B promoter.
- Circuit A produces the proteases that degrade the proteins produced by circuit B.

Furthermore, in each scenario we considered two different configurations for circuit A (open loop and negative feed-back regulated), and for each configuration we varied the degradation rate of circuit B proteins.

Our results indicate that, the quasi-oscillatory behavior of circuit B is quite robust to random variations of the transcription factor count, independently of the fluctuations time scale, and of whether this suorce of extrinsic noise is quasi-periodic or correspons to colored noise.

Contrarily, we found that the quasi-periodic behavior of circuit B deteriorates when the degradation rate of circuit B proteins randomly varies in time due to extrinsic noise. Moreover, the extrinsic noise effects become more notorious as the time scale of the corresponding fluctuations gets shorter.

Summarizing, we investigated the effects of extrinsic noise on the quasi-periodic behavior of a gene network subject to delayed negative-feedback regulation. According to our results, these effects strongly depend on the parameters affected by extrinsic noise, as well as on the time scale of the noisy fluctuations. In most cases we observed that low frequency extrinsic noise has little effect upon the system oscillatory dynamics.

## Acknowledgments

The author is thankful to Prof. E. S. Zeron for his help and advice in the development of the present project.

## Reference

[1] A. Goldbeter, Biochemical Oscillations and Cellular Rhythms: The Molecular Bases of Periodic and Chaotic Behaviour (Cambridge University Press, 1997).

[2] G. Bordyugov, P. O. Westermark, A. Korenffčič, S. Bernard, and H. Herzel, in Circadian Clocks (Springer Science, 2013), pp. 335–357.

[3] R. Kageyama, Y. Niwa, A. Isomura, A. González, and Y. Harima, WIREs Dev Biol 1, 629 (2012).

[4] O. Purcell, N. J. Savery, C. S. Grierson, and M. di Bernardo, Journal of The Royal Society Interface 7, 1503 (2010).

[5] V. Singh, Gene 535, 1 (2014).

[6] I. Lestas, J. Paulsson, N. E. Ross, and G. Vinnicombe, IEEE Transactions on Automatic Control 53, 189 (2008).

[7] R. Silva-Rocha and V. de Lorenzo, Annu. Rev. Microbiol. 64, 257 (2010).

[8] V. Shahrezaei, J. F. Ollivier, and P. S. Swain, Mol Syst Biol 4 (2008).

[9] H. Ge, H. Qian, and X. S. Xie, Phys. Rev. Lett. 114 (2015).

[10] N. MacDonald, Biological Delay Systems: Linear Stability Theory (Cambridge Studies in Mathematical Biology) (Cambridge University Press, 1989).

[11] J. Lewis, Current Biology 13, 1398 (2003).

[12] B. Novák and J. J. Tyson, Nature Reviews Molecular Cell Biology 9, 981 (2008).

[13] J. Wang, M. Lefranc, and Q. Thommen, Biophysical Journal 107, 2403 (2014).

[14] A. Sanchez and I. Golding, Science 342, 1188 (2013).

[15] D. Kennell and H. Riezman, Journal of Molecular Biology 114, 1 (1977).

[16] T. Erneux, *Applied Delay Differential Equations* (Springer New York, New York, 2009).

[17] J. M. Raser, Science 309, 2010 (2005).

[18] http://www.computerhistory.org/revolution/analog-computers

[19] http://www.arduino.cc/

[20] http://www.bitscope.com

